# Stochastic population dynamics driven by mutant interactors

**DOI:** 10.1101/397810

**Authors:** Hye Jin Park, Yuriy Pichugin, Weini Huang, Arne Traulsen

## Abstract

Spontaneous random mutations are an important source of variation in populations. Many evolutionary models consider mutants with a fixed fitness chosen from a certain fitness distribution without considering any interactions among the residents and mutants. Here, we go beyond this and consider “mutant interactors”, which lead to new interactions between the residents and invading mutants that can affect the carrying capacity and the extinction risk of populations. We model microscopic interactions between individuals by using a dynamical payoff matrix and analyze the stochastic dynamics of such populations. New interactions drawn from invading mutants can drive the population away from the previous equilibrium, and lead to changes in the population size — the population size is an evolving property rather than a fixed number or externally controlled variable. We present analytical results for the average population size over time and quantify the extinction risk of the population by the mean time to extinction.

## I. INTRODUCTION

Evolutionary game theory has been widely used to describe the evolution of populations under frequency dependent selection [1, 2]. Conventionally, most evolutionary game theoretical models are based on fixed or infinite population size and thus only reflect changes in the frequencies of types. As an increasing body of research has shown that evolutionary and ecological processes can happen on comparable time scales [3–10], considering both the evolutionary processes arising from mutations and the ecological effects due to an associated change of the total population size is important [12–20]. Most eco-evolutionary dynamics has been explored theoretically based on deterministic equations such as the competitive Lotka-Volterra equation [21] or extensions of the replicator dynamics [22]. These approaches describe the changes of abundances in time, but they cannot naturally describe possible stochastic events such as a population extinction. Instead, individual-level models can capture the stochastic nature of the population dynamics better by using microscopic approaches based on reaction rules [20, 23–25]. Although stochastic systems driven by such microscopic reaction rules have been traditionally studied in mathematical biology [26], the stochastic dynamics of an evolving population driven by mutants has not been studied so far.

Based on a game theoretical approach with a dynamic payoff matrix [11], we consider both mutations leading to new interactions and the changes in the population size caused by those interacting types; Mutations and extinctions of types are captured by the extension or reduction of the payoff matrix, where the new entries related to the mutant are randomly drawn from the corresponding entries of the maternal type [11]. By introducing death through competition between individuals [20], the population is naturally controlled by individual interactions and evolves over time. This framework provides us a general framework to study the long term evolution of interacting populations through the analysis of present interactions.

In this manuscript, we especially focus on the evolution of the population size over time, as it has an important impact on the eco-evolutionary dynamics. The emergence of a new mutant can lead to an increase or a decrease of the population size depending on the new interactions between residents and mutants. If a mutant type with a smaller payoff outcompetes the resident type with a higher payoff, a so-called *s*ocial dilemma situation, the population size decreases. If the population size consecutively decreases due to the invasion of such mutants, the population may even go extinct. We analyze the changes of the carrying capacity, which is defined as the average population size when the population composition reaches a stable state. Thus, the carrying capacity is an evolving rather than a predefined property of our stochastic processes.

We also obtain the long time behavior of the population size and estimate the mean time to extinction by mapping the problem to a random walk. Counterintuitively, we find that the carrying capacity and the mean time to extinction do not monotonically increase with the probability *θ* that a new payoff for the mutant is larger than the maternal payoff. Especially, for small probability *θ* there is a tradeoff between a large decrease in the population size and a small chance of such a mutant reaching fixation.

This manuscript is structured in the following way: we introduce our model in detail in Sec. II and present an analysis of the properties of the model in Sec. III. First, we calculate the population size changes induced by one mutation event in a single-type population. Then, we assess the evolution of the carrying capacity in the long run and estimate the extinction risk based on the mean time to extinction. Finally, we summarize and discuss our results in Sec. IV.

## II. MODEL

We study the stochastic population dynamics with a mutation process that constantly creates new types in the population. A mutation occurs with probability *μ* during reproduction. Reproduction and intrinsic death occur at constant rates, λ_*b*_ and λ_*d*_, respectively (λ_*b*_ > λ_*d*_). Individuals also die due to the competition for a limited resource. These competitive interactions are modelled based on a game theoretical approach, where the death rates from competition are the inverse of individual’s payoffs. All these microscopic reaction rules are summarized in Table I.

**TABLE I.**
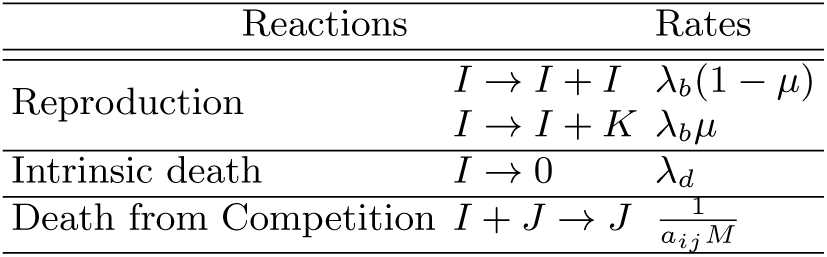
Reaction rules with corresponding rates. The parameter *M* controls the scale of the population size. The payoff of a type *i* interacting with a type *j* is denoted as *a*_*ij*_. Mutation occurs with probability *μ;* during reproduction and leads to an extended payoff matrix.

For large populations without mutations, a deterministic equation describes the change of the abundance of type *i, x*_*i*_,

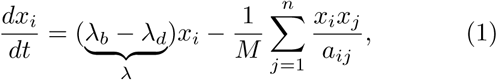

where *n* is the current number of types in the population and *a*_*ij*_ is the payoff of a type *i* from the interaction with a type *j*. When the population size is of the order of *M*, the reproduction and death occur at similar rates, such that the population neither grows nor shrinks. Accordingly, the population size scales in *M*. For single-type populations with the single payoff *a*, population size is fluctuating around the fixed point *K* = *aM*λ. The population size *K* at the stable fixed point of Eq. (1) is considered as the carrying capacity of the population.

We assume that mutations occur sufficiently rarely. The equilibration time of the population composition after the emergence of a mutant is much shorter than the waiting time between consecutive mutations, see Fig 1. Thus, the population either stays close to one of the monomorphic populations or becomes polymorphic and fluctuates around a coexistence equilibrium in between two successive mutation events. The population size between two successive mutation events is characterized by the carrying capacity calculated from interactions between individuals. We use a discretized time *t* based on mutation events. Once a new mutant emerges, the mutant event time *t* increases by one. Then, we use *K*, the carrying capacity at the equilibrium, as the characteristic population size.

**FIG. 1.**
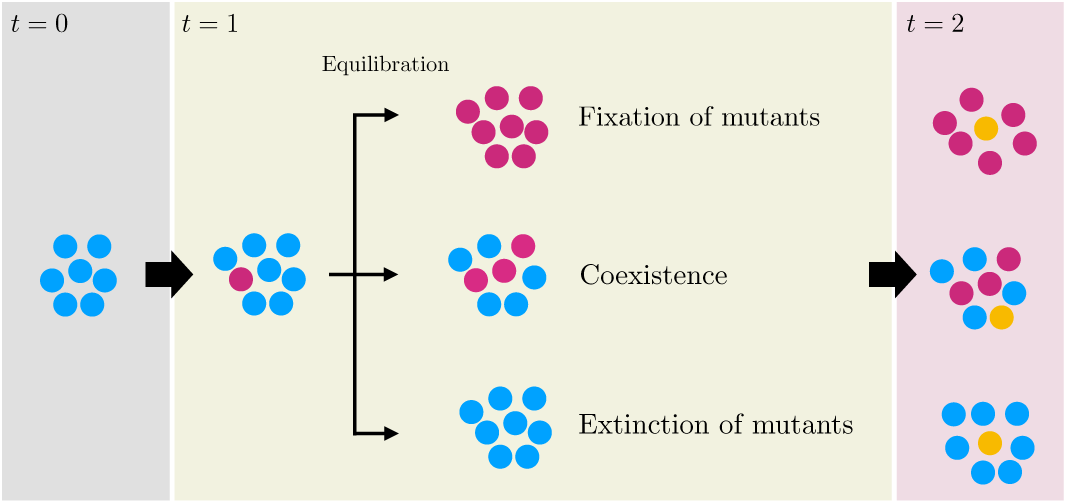
There are three possible outcomes from a mutation event: (i) the mutant spreads through the population and replaces the residents, (ii) the mutants coexist with the residents, or (iii) the mutants go extinct. When the waiting time for the emergence of the next mutant is long, the population has enough time to reach one of these outcomes. After equilibration, the population size fluctuates around the new carrying capacity before the next new mutation event. Hence, the population size between two successive mutation events is characterized by the carrying capacity calculated from interactions between individuals. We discretize time by mutation events and use the carrying capacity as a characteristic population size in the weak mutation regime. While most of our analytical results are carried out under the weak mutation assumption, equilibration before the next mutation is not necessary for stochastic simulations. Details for the stochastic simulations are available in Appendix A.

Once a mutant emerges in a population with *n* types, the payoff matrix extends its size from *n* × *n* to (*n* + 1) × (*n* + 1). For example, if the mutant emerges in a single-type population with payoff *a*_11_, we can write down the change of the payoff matrix as

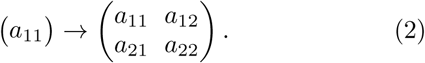

The payoff *a*_11_ is for the interaction between residents, and *a*_22_ is for the interaction between mutants. The payoffs *a*_12_ and *a*_21_ are for the interaction between a resident and s mutant. New payoffs are randomly drawn from a distribution controlled by the probability *θ* that a new payoff is larger than the maternal payoff. In principal, we can use any distribution here [27], but for simplicity we focus on the exponential distribution. In the example Eq. (2), there is only one maternal payoff *a*_11_, and hence, new payoffs *a*_12_, *a*_21_ and *a*_22_ are independently drawn from the same distribution 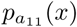 given by

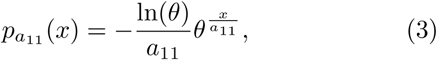

which satisfies 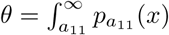.

When a mutant emerges in populations with several types (*n* > 1), it is denoted as the (*n* + 1)th type. The payoffs *a*_*i, n*+1_ of a resident type *i* from the interaction with this mutant type are based on the payoffs *a*_*im*_ from the interaction of the same resident type with the maternal type *m* [11]. In other words, the last row entries *a*_*i, n*+1_ are drawn from the distribution 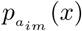 and the last column entries *a*_*n*+1*, i*_ are drawn from 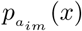. The emergence of a mutant may lead to a change in an equilibrium, and the population size fluctuates around a new carrying capacity if the mutant establishes itself in the population.

## III. RESULTS

Each time a mutant emerges in the population, the mutation event time *t* increases by one, and either new mutants establish themselves in the population or they die out, see Fig. 2. Due to the new interactions from mutants, fixation of mutants or coexistence between mutants and residents induce a change of the carrying capacity. Here, we focus on the short and long time dynamics of the carrying capacity, which is induced by such new interactions. Since the probability *θ* is directly connected to the new payoffs, we use *θ* as a key control parameter. For the short time evolution, we look at the average carrying capacity changes from the emergence of a single mutant type in III A. For the long time evolution, we get asymptotic behavior of the carrying capacity and calculate the mean time to extinction in III B.

**FIG. 2.**
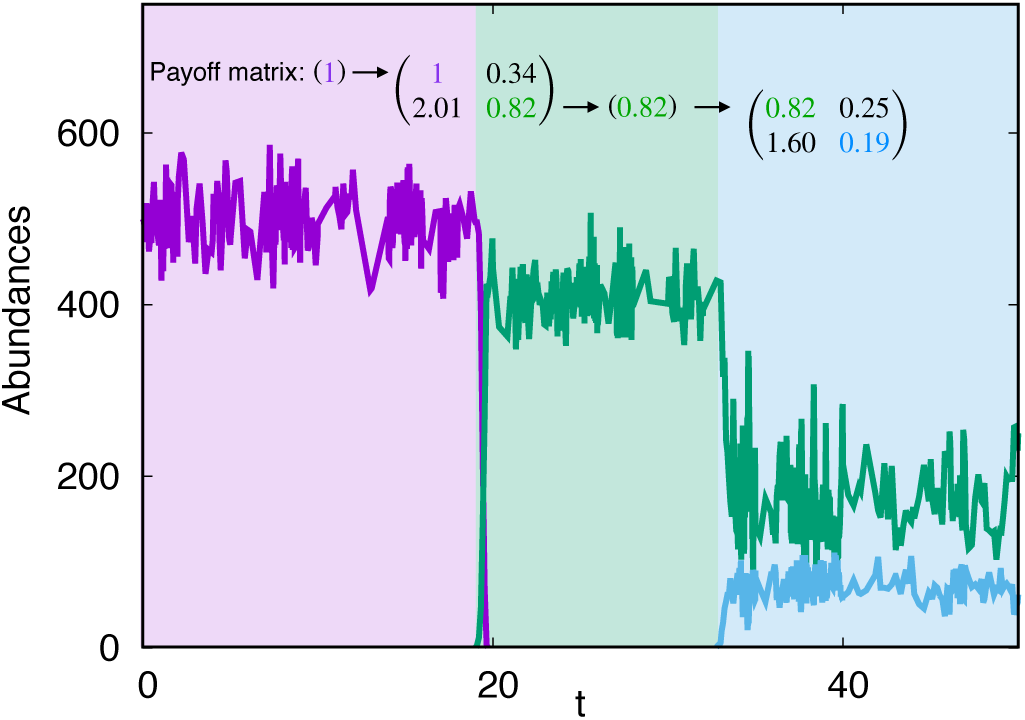
One realization of the stochastic simulation showing abundances *x*_*i*_ in mutation event time *t*. We measure abundances in real time *T* and then convert *T* to *t* making one mutation event increase *t* by one. We use a single-type population as an initial condition. Lines of different colors indicate abundances of different types. Here, we only shown successful mutants which establish themselves in the population. The abundances *x*_*i*_ fluctuate around the equilibrium before the successful invasion. There are three possible situations after a mutant emerges from a single-type population. (1) mutants die out, as in the majority of shown events. (2) mutants take over the whole population, as the green type emerging at *t* = 19. (3) mutants and residents coexist, as the green and cyan types after *t* = 32. We used *a*_11_ = 1, *M* = 1000, λ_*b*_ = 0.9, λ_*d*_ = 0.4, *θ* = 0.2, and *μ* = 5 × 10^−5^.

### A. Changes of carrying capacity induced by a single mutation event

In a large population containing a single resident type with the payoff *a*_11_, the population size fluctuates around *K* = *a*_11_*M*λ. Once a mutant emerges, new equilibria may arise depending on the payoffs, see Eq. (4). For the possible new carrying capacities, we look at the stability of new equilibria. For *a*_21_ < *a*_11_, a mutant receive smaller payoff than the resident, while the mutant is rare. Thus, either the resident type dominates the mutant type (*a*_12_ > *a*_22_, resident dominance game) or both homogeneous populations are stable (*a*_21_ < *a*_22_, coordination game). In these two cases, a mutant typically gets lost. For *a*_21_ > *a*_11_, the mutant type is likely to invade from rare. Here, the mutant either dominates the resident type (*a*_12_ < *a*_22_, mutant dominance game) or they coexist (*a*_12_ > *a*_22_, coexistence game), and a new equilibrium is achieved [20]. In summary, if the population successfully reaches their new stable equilibrium, the new carrying capacity after the mutation becomes

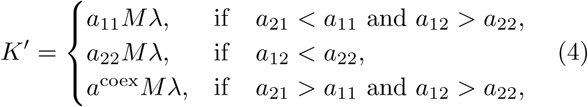

where 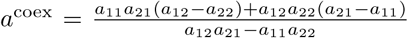 is the average payoff in the coexistence equilibrium.

Note that these equilibria are not guaranteed to be reached. In a stochastic process, the abundance of the mutant type can have large random fluctuations especially at the beginning because it starts from a single individual. These fluctuations can lead to the extinction of the mutant type even if it has dominating payoffs (*a*_21_ > *a*_11_ and *a*_22_ > *a*_12_). Therefore, to calculate the carrying capacity changes, we have to estimate the probability *ϕ* of a mutant to successfully establish itself in the population. Once *ϕ* is known, the average change of the carrying capacity is given by

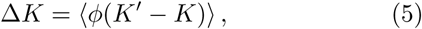

where *K* = *a*_11_*M*λ and bracket ⟨*…*⟩ represents the averaging over all mutant types which are distinguished by a different payoff matrix.

The risk of stochastic extinction becomes negligible when the mutant type reaches a large abundance. Thus, *ϕ* is determined mainly in the early stage of an invasion. For *a*_21_ > *a*_11_, the extinction risk of mutants is quickly reduced with increasing abundance of the mutants, *y*. We assume that the mutants successfully escape from stochastic extinction and settle in the population if their abundance reaches *y*^*\**^, 1 ≪ *y*^*\**^≪ *M*. During this change, we also assume that the total population size and the abundance *x* of the resident type do not change significantly, *x* ≈ *a*_11_*M*λ = 𝒪 (*M*). Therefore, we use the fixation probability of mutants in a population with a constant size to estimate *ϕ* as [28, 29],

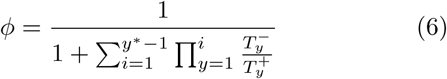

where 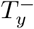 and 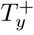 are the rates of decrease or increase the number of mutants by one, starting in *y*. For large *M* and small *y, y* ≪ *M*, these can be approximated by

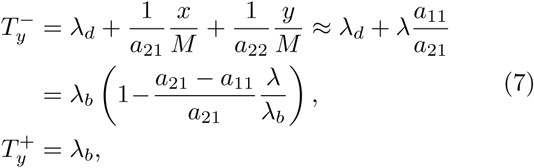

such that 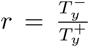 becomes independent of *y* and constant. From this, we obtain an approximated expression of the probability *ϕ* for *a*_21_ > *a*_11_,

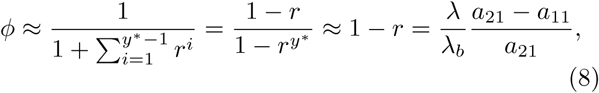

where in the last step we took into account that *y*^*\**^ ≫ 1 and *r* < 1.

For *a*_21_ < *a*_11_ and *a*_12_ > *a*_22_, the homogeneous resident population is stable against invasions, and the mutant cannot settle in the population. However, for *a*_21_ < *a*_11_ and *a*_12_ < *a*_22_, both single-type populations are stable and have their own basins of attraction [30, 31]. Taking a closer look at Eq. (1) and its vector field, we find that when *a*_12_ is sufficiently small, an emergence of a single mutant is enough to put the population into the basin of attraction of another equilibrium – the homogeneous mutant population.

In coordination games, 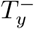 is greater than 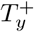 at the beginning (*x* ≈ *a*_11_*M*λ and *y* ≈ 1), so the mutant population *y* is more likely to shrink rather than increase in numbers. If *a*_12_ is small enough, the same is true for the resident population and the abundance of residents will decrease much faster than the abundance of mutants. Once *x* becomes smaller than *a*_21_*M*λ, 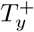 becomes larger than 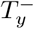, and after this point the mutant population begins to grow. Hence, if the mutants survive until the abundance of residents *x* decreases to *a*_21_*M*λ, they will fixate in the population.

The probability that the single mutant does not die during the time *t*_*s*_, while *x* decreases from *a*_11_*M*λ to *a*_21_*M*λ, can be approximated by 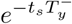. We calculate the time *t*_*s*_ by looking at the rates of increasing or decreasing the abundance of the resident type,

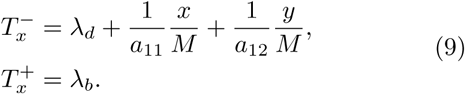

When *a*_12_ is very small, *a*_12_ = 𝒪 (1*/M*), the third term in 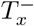 is not negligible even if *y* is small. Given the large initial abundance of the resident type, its dynamics can be described by the deterministic Eq. (1). Then, we get the decrease rate of the resident’s population size

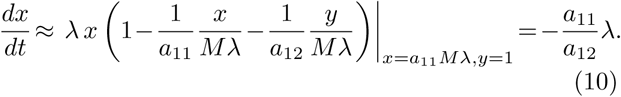

From the estimation, we calculate the time *t*_*s*_ it takes from 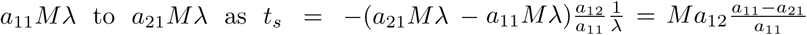. Thus, the fixation probability of mutants in a coordination game can be approximated by

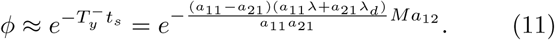

Note that although these mutants have a very small chance to reach fixation in the population, the coordination game (*a*_21_ < *a*_11_ and *a*_12_ < *a*_22_) plays an important role for small *θ*. In this regime, the fixation of dominant mutants in the population is becoming very unlikely – instead fixation in coordination games becomes the dominant factor of evolution. Most importantly, this small probability strongly influences the long time behavior for small *θ*. In summary, the probability *ϕ* can be approximated as

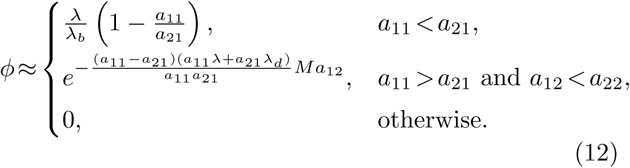

As the probability *ϕ* of the mutant in coordination games becomes relevant only if *a*_12_ ∼ 1*/M*, we only consider mutant dominance and coexistence games in calculating Δ*K* for the short time behavior. Hereafter, we refer to the mutant dominance game as dominance game because the resident dominance game does not play a role for changing the carrying capacity. In Fig. 3 (a), we numerically show that an invasion of a coexisting mutant leads to much smaller changes in the carrying capacity than a fixation of a dominant mutant without stochasticity. Hence, we make a further approximation assuming that only dominant mutants contribute to the change of the carrying capacity. Under this approximation, the change in the carrying capacity after a single mutation event can be calculated analytically

**FIG. 3.**
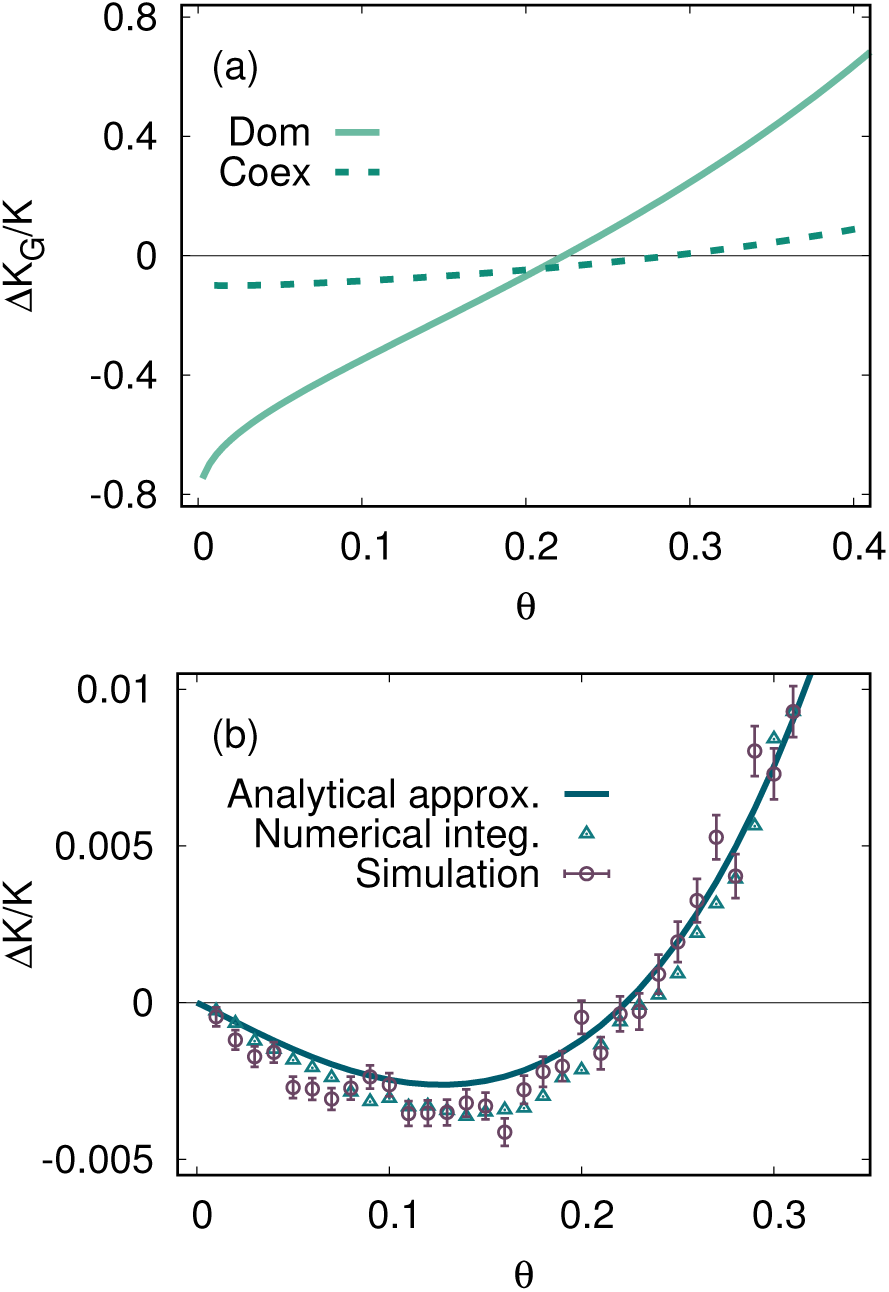
Relative changes in carrying capacity induced by a single mutation event. (a) the changes Δ*K*_*G*_ of relative carrying capacity without stochasticity at a given game type are shown, in the dominance game (solid line) or the coexistence game (dashed line). The changes induced by coexisting mutants are much smaller in magnitude than changes from dominant mutants. (b) the average change of the carrying capacity after a single mutation event in stochastic simulations (brown circles), a numerical integration of Eq. (5) (green triangles), and our analytical approximation Eq. (13) (solid line) are shown. The results of stochastic simulations are well matched by our approximation. We used an initial payoff *a*_11_ = 1, *M* = 1000, λ_*b*_ = 0.9, and λ_*d*_ = 0.4. For stochastic simulations, we used 50000 realizations.

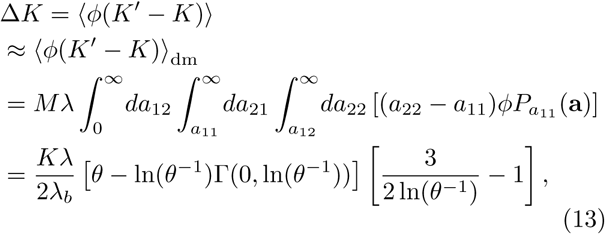

where subscript “dm” indicates that the averaging is performed only across dominance games, and 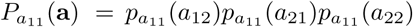 with **a** = (*a*_12_*, a*_21_*, a*_22_). The incomplete Gamma function is given by 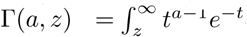. Our stochastic simulations agree with this approximation, see Fig. 3 (b).

For a given game type, once mutants that decrease the population size settle in the population, the smaller *θ*, the larger the drop of the population size as shown in Fig. 3 (a). Interestingly, however, the change of the relative average carrying capacity of mutants does not monotonically increase with *θ*, see Fig. 3 (b). Smaller *θ* induces larger decreases of the population size, but these changes also become exceedingly rare. From those two effects of small *θ*, we observe a large drop of an average population size for intermediate *θ*.

### B. Evolution of the carrying capacity in the long run

In the previous subsection, we have studied the carrying capacity changes induced by a single mutation event. Coexistence games do not change the population carrying capacity *K* much, while the fixation of the mutants more dramatically change *K*. Usually, the change of *K* from the coexistence game is negligible and thus we only consider fixation of mutants in our analytical calculation for the long time behavior. As a consequence, the dynamics of the population can be approximated as a sequence of single payoffs which induce changes in the population size. In this subsection, we map our problem to a random walk in the payoff space to obtain the evolution of the carrying capacity and approximate the mean time to extinction.

If a new mutant type fixates in the population, *a*_22_ becomes the new single payoff. The distribution of *a*_22_ as a new single payoff in the dominance game is given by

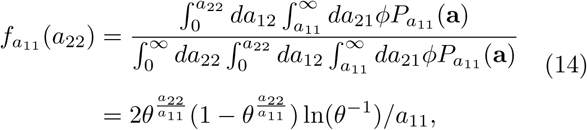

which depends on both *a*_11_ and *a*_22_. By defining *l* = ln 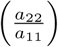, we can convert the distribution 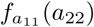 into a jump distribution *f* (*l*), which is a function of a single variable,

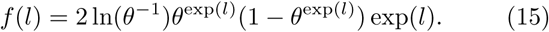

We define *a*^(*τ*)^ as the payoff of *τ*-th successful fixating mutant type. After every successful fixation event, the logarithm of the payoff jumps by a distance *l*, ln (*a*^(*τ+*1)^) = ln (*a*^(*τ*)^) +*l*. Hence, the evolution of the logarithm of payoff maps into the random walk problem with the jump distribution Eq. (15). When the context is clear, we omit the superscript *τ*. Once the population size goes down (*a* is decreasing), fixations from mutants playing coordination games become more relevant because a chance to get a small enough *a*_12_ increases. Hence, we also consider the coordination game as well as the dominance game in the long time evolution. The jump distributions derived from dominance and coordination games become more similar as *a*_11_ decreases, so Eq. (15) continues to be applicable for the coordination game.

For the sake of simplicity, we use the fixation time *τ*, a unit of the single fixation event, and treat it as continuous variable. Then, we can write down the diffusion equation to describe the dynamics of *u*(*τ*) = ln(*a*^(*τ*)^)

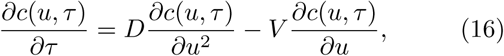

where *c*(*u, τ*) is the probability density of a random walker at position *u* at time *τ*. The diffusion coefficient *D* and the drift velocity *V* are given by the moments of distribution (15), 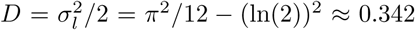 and *V* = ⟨ *l* ⟩ = −*γ* + ln(2) − ln(ln(*θ*^−1^)) where *γ* ≈ 0.544 is Euler’s constant.

The boundary condition to this equation is given by the stochastic extinctions of population. Since the population is likely to go extinct when the population size is small, there is a threshold payoff which can be defined as an absorbing boundary. Based on simulation results, we measure the threshold payoff *ã*(*θ*) under which populations go extinct before the next fixation event (see Appendix D). Under this boundary condition *c*(ln(*ã*)*, τ*) = 0 and the initial condition *c*(*u*, 0) = *δ*(*u*, 0), the solution of Eq. (16) yields (see [32], p. 87),

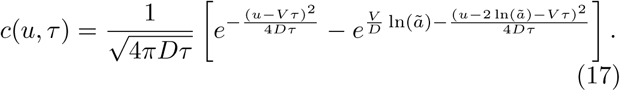

Using this solution, we obtain the expectation value of the carrying capacity in the long run (see Appendix E for details)

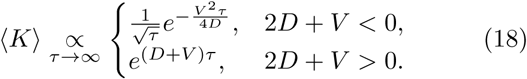

The expected value increases for *D* + *V* > 0 (*θ* > 0.206) and decreases for *D* + *V* < 0 (*θ* < 0.206).

If the population size increases, it eventually becomes too large to satisfy the weak mutation assumption. The large population size decreases the time interval between consecutive mutation events and increases the time to equilibration. In this case, a new mutant emerges before the equilibration of the population. Thus, the population size does not reach the new carrying capacity calculated from the payoff matrix, and the population size *N* cannot be characterized by the carrying capacity anymore. Since the interactions play the role if the population size is close to the carrying capacity, the competition becomes negligible and the population grows at a maximum rate of the order of λ, see Eq (1). The average population growth rates in this regime agree well with our prediction, as shown in Fig. 4, see Appendix F for details. Note that the average population size ⟨ *N* ⟩ does not always increase for *θ* > 0.206 in simulations. This may arise from the emergence of mutants who play coexistence games with resident. The changes of population size induced from dominance games are much larger than that of coexistence games, except in the vicinity of *θ* which induces zero change of the population size in dominance games, *θ* ≈ 0.223 (see Fig. 3). As mutants who play coexistence games decrease the population size for *θ* > 0.206 while dominance games induce very small changes in the population size as shown in Fig. 3 (a), the average population sizes are reduced even for *θ* > 0.206.

**FIG. 4.**
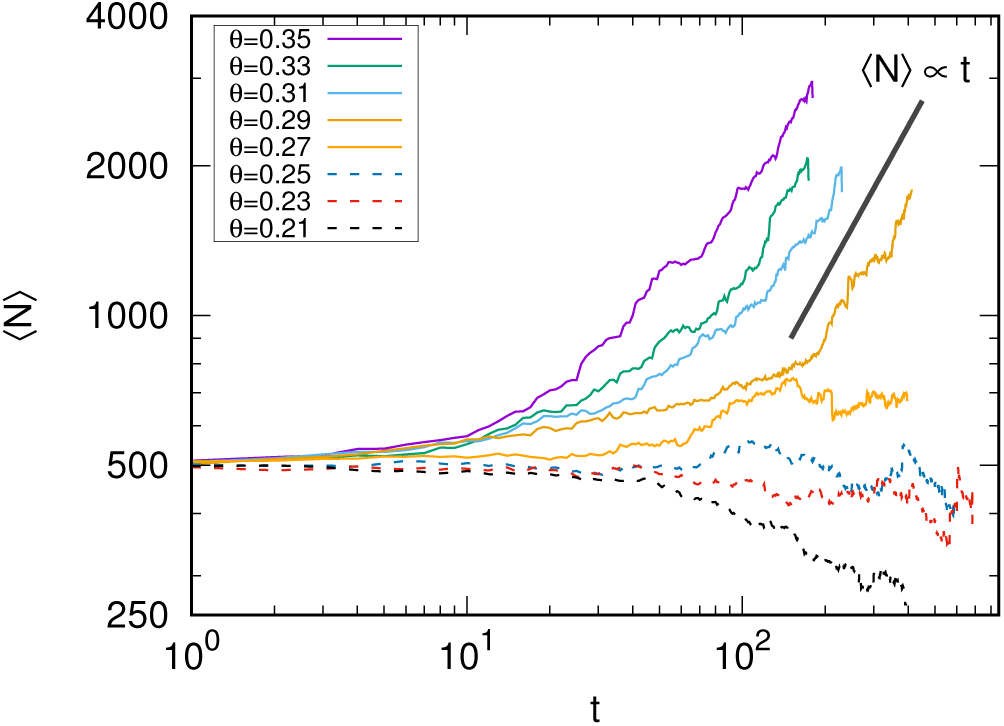
Simulation results of the average population size ⟨ *N* ⟩ in mutation events *t* for *θ* > 0.206. We used dashed and solid lines for showing trajectories where the the population sizes decrease or increase at the end of the simulations. We use log scales for both axes, and the thick black solid line is a linear function in time *t* for comparing the trend of population size changes and the linear function. As we can see, the asymptotic behavior of the population growth is linear in time *t* which is expected out of the weak mutation regime, see Appendix F for details. We used an initial payoff *a*_11_ = 1, *M* = 1000, λ_*b*_ = 0.9, λ_*d*_ = 0.4, and *μ* = 10^−5^ (500 realizations for each *θ*).

If the population size decreases, the population will eventually go extinct. However, the time to reach the extinction differs for different *θ*. We calculate the mean time to extinction, *t*_ext_, for *θ* < 0.206 to estimate the extinction risk. The extinction rate *k*_ext_(*τ*) at *τ* is the density current passing through the absorbing state (*u* = ln(*ã*)),

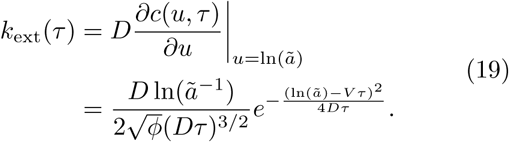

Hence, we can get the mean time to extinction *τ*_ext_

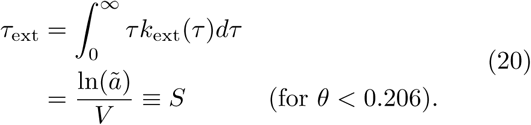

This is exactly the same as the necessary time to move the distance ln(*ã*) with velocity *V*. This result is based on the fixation time unit, and thus *S* stands for the expected number of random walker’s jumps before extinction.

The random walker jumps only if the mutants successfully fixate in the population, and not every mutation leads to a successful fixation. Now, we find the expected number of mutations before extinction. The jump rate *ξ* of the random walker is determined by the combined probabilities *ϕ* for dominance and coordination games

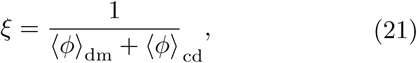

where

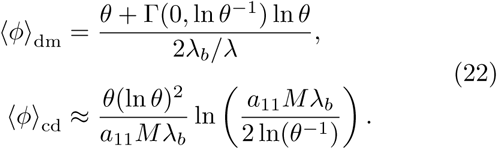

The probability ⟨ *ϕ* ⟩_dm_ only depends on *θ* while ⟨ *ϕ* ⟩_cd_ is a function of both *θ* and the payoff *a*_11_ (see Appendix B for the exact expression). On average, *ξ* mutations happen until one successfully fixates.

As *ξ* mutation events are needed on average for each jump, the mean time to extinction in the unit of *t* is

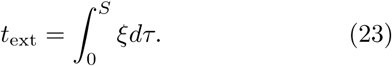

We use the relation ln(*a*^(*τ*)^) = ln(*a*^(0)^) + *V τ* and calculate the mean number of mutations before extinction as follows,

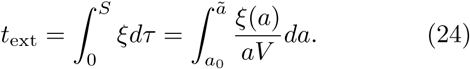

The expression for *ξ* is available in Eqs. (21-22). We numerically compute *t*_ext_ according to Eq. (24) and compare the results with stochastic simulations in Fig. 5. Our analytical results well predict the simulation outcomes. Interestingly, again the mean time to extinction does not monotonically increase with *θ*. As the same with the short time changes of the carrying capacity, a minimum *t*_ext_ is observed in the intermediate *θ* due to the small *θ* properties. Smaller *θ* implies a larger decrease of the population size. At the same time, small *θ* leads to long waiting time for changes in the carrying capacity. Hence, there is a trade-off between large jumps to the small population size and long waiting time for such a jump.

**FIG. 5.**
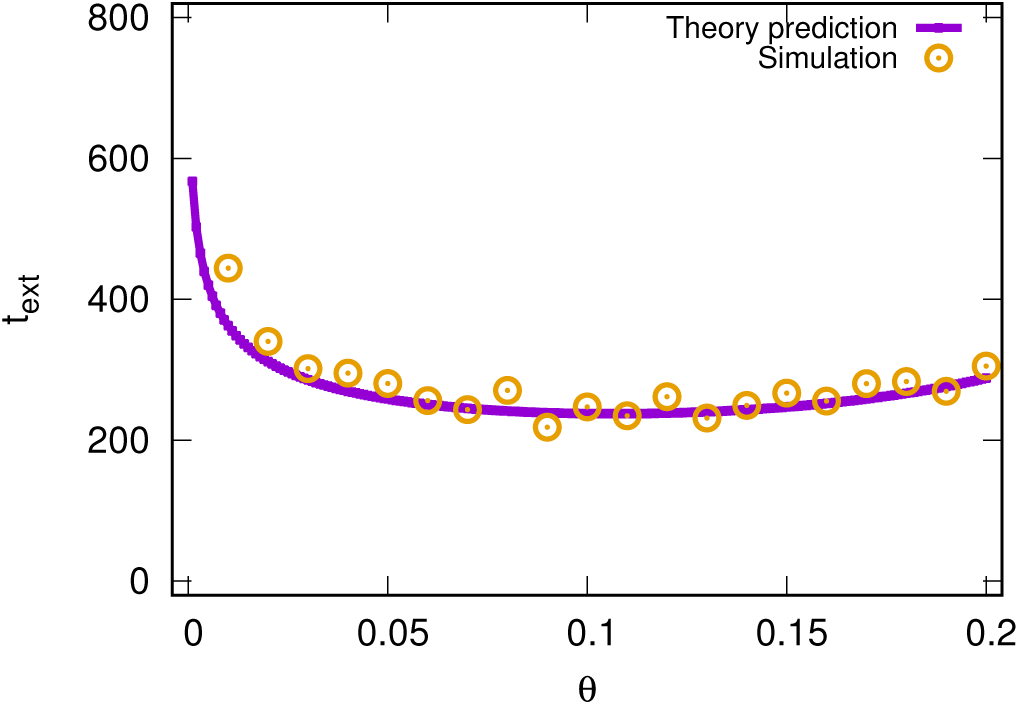
The mean time to extinction *t*_ext_ in *θ*. Stochastic simulation results are shown as open circles. The solid line is our theoretical prediction (24). Similar to the short time behaviour, the mean time to extinction also shows a minimum at the intermediate *θ* due to the small *θ* properties. Smaller *θ* induces larger decreases of the population size, while it also delays such decreases. Our analytical approximation predicts the mean time to extinction well. We used *M* = 1000, λ_*b*_ = 0.9, λ_*d*_ = 0.4, and *μ* = 10^−5^ with an initial payoff *a*_11_ = 1.

## IV. SUMMARY AND DISCUSSION

We implement a stochastic model with mutations leading to novel interactions and changes of the population size. We focus on the evolution of the population size and the mean time to extinction. Since we interpret competition as a game theoretical interaction, the relation between types and thus the population size are determined by payoffs. In a social dilemma, the population size decreases if a mutant type outcompetes the resident type – but the mutants have stronger competition among themselves. This phenomenon is similar to mutational meltdown from deleterious mutations [33–35]. In our model, the distribution of new payoffs is important, because it controls how often such deleterious mutations happen. If the probability *θ* to have a larger payoff than the maternal payoff is too small, populations are endangered by deleterious mutations. On the contrary, when *θ* is large enough, the population size constantly increases. For short and long time evolution, we find a trade-off between large decreases of the population size and the rareness of such events.

Although the model is governed by simple reaction rules, intriguing phenomena emerge. The investigation of different distributions of new payoffs and its correlation is a challenging problem to be addressed in future work. While we only focus on monomorphic populations in the main text, our model can be used also to investigate polymorphisms [11].

In long-term experiments of microorganisms, parallel mutations are often seen in populations derived from a common ancestor [36–41]. By tracking the point mutations in single nucleosides over thousands of generations, e.g. in the seminal experiments by Lenski et al., rich population dynamics including selection sweeps by mutations with fitness advantages, are observed [42], but the experimental conditions preclude loss of the population. For this, a more complex setup is necessary [43]. Also quasistable coexistences of multiple types have been observed in recent experiments [40, 41].

Our model provides a general framework to model general eco-evolutionary processes in natural populations, as it has no artificial restriction on the population dynamics, i.e., the population size and composition evolve solely depending on random mutations and simple dynamic rules. We hope that this model will inspire other researchers to work on evolutionary models in which interactions are not pre-defined but evolve *de novo* with a natural link to demographic fluctuations.

## APPENDICES

### Appendix A: Stochastic simulations

To implement the reaction rules in Table I, we use an algorithm developed in [44] which is similar to the Gillespie algorithm [45]. In the algorithm, one of the reaction rules is attempted at a given small enough time interval, and the simulation time proceeds based on this try leading continuous time. We call this simulation time the real time *T* and can trace changes in the population size in *T*. When the mutation rate is small enough, however, the population size is fluctuating around the carrying capacity determined by Eq. (1). Then, almost all simulation time after equilibration of the population does not contain new information. Hence, we use a discretized time *t* based on the mutation events and only take the population size right before the emergence of the new mutant type as the representative population size at a given *t*. In general, the waiting time for the new mutant type is long enough when the mutation rate is small, and thus the carrying capacity can be representative for the population size. However, according to the reaction rates, a new mutant type can arise before the equilibration of the population. In this case, the carrying capacity may not represent typical population size. Core codes are available at https://github.com/Park-HyeJin/MutantInteractors.

### Appendix B: **The exact expression of *ξ***

The probability ⟨*ϕ*⟩ _dm_ that a mutation will result in a fixation of dominating mutant is

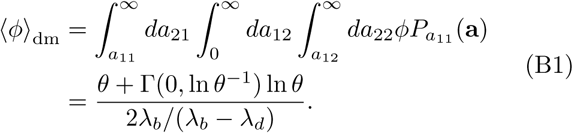

The probability does not depend on the resident payoff value *a*_11_ while the contribution ⟨*ϕ*⟩_cd_ from the coordination game depends on *a*_11_ as well as *θ*. The probability *ϕ* integrated in coordination game is

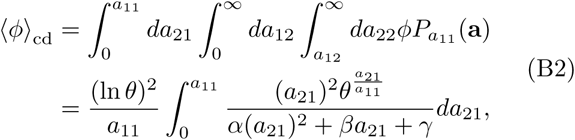

where *α* = *a*_11_*M*λ − *a*_11_*M*λ_*b*_ − 2 ln *θ*, 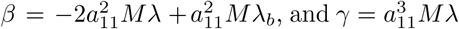. For our parameters of interest, the above integration can be solved,

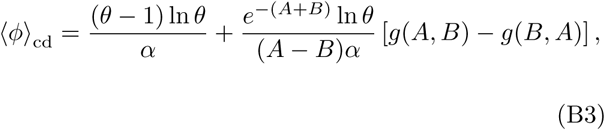

where 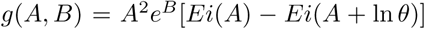 with 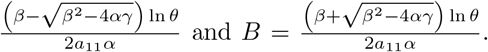. The function *Ei*(*z*) is the exponential integral function 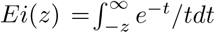. Thus, the average probability is depending on the current payoff *a*_11_.

Furthermore, we get the approximated expression of the probability ⟨*ϕ*⟩ _cd_ from Eq. (B2), which leads to the integral

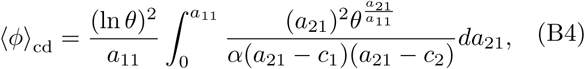

where

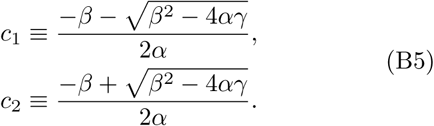

For large *M*, we obtain the approximated expressions

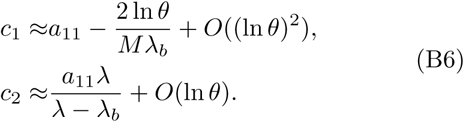

Note that the function inside of the integration diverges at *a*_21_ = *c*_1_ and *a*_21_ = *c*_2_. Since *c*_1_ is close to *a*_11_, an upper limit of integration, the major contribution of the integration comes from the vicinity of this upper limit. We utilize this diverging behavior to simplify the expression of Eq. (B4). We do a partial fraction decomposition and neglect non-diverging term

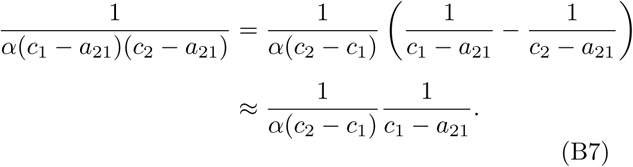

**FIG. B1.**
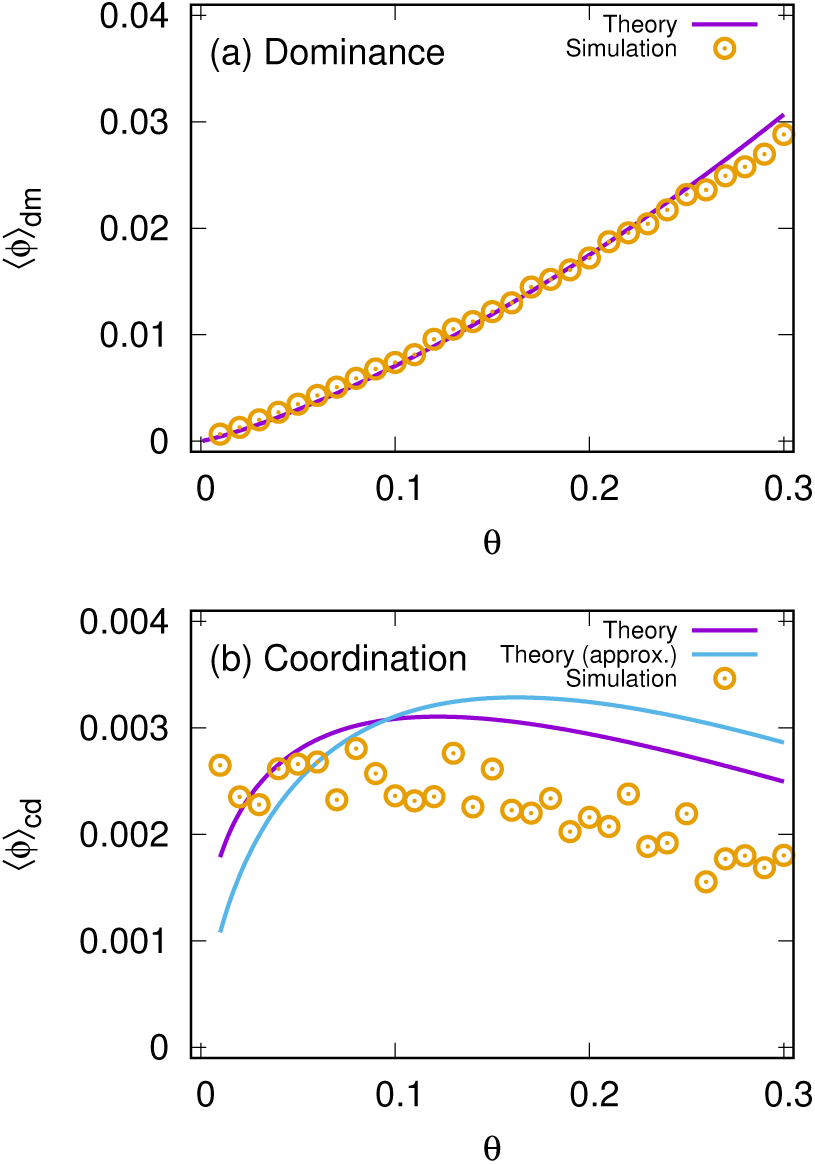
We show the average settling probability ⟨*ϕ*⟩ for (a) dominance game and (b) coordination game. The theoretical prediction Eqs. (B1) and (B3) are shown as purple lines. For the coordination game, we show the approximation given by Eq. (B10) (solid line). For stochastic simulations, we use 20000 trials of the dominance or coordination games with *a*_11_ = 1 in each point. Especially, the probability in the dominance game agrees well with the theory. For the coordination game, our approximation does not work well but reproduces the right trend in the right order of magnitude. We used *M* = 1000, λ_*b*_ = 0.9, and λ_*d*_ = 0.4.

Similarly, we approximate the nominator in integration as 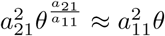. Hence, we get the expression

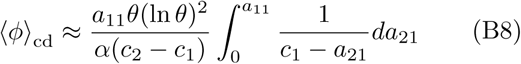

The prefactor can be approximated by

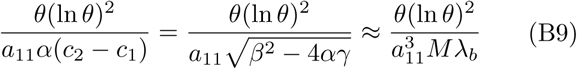

for large *M*. Finally, we get

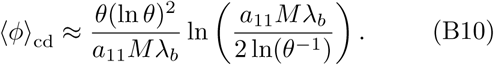

We compare the stochastic simulation results and theoretical prediction for *a*_11_ = 1 in Fig. B1.

### Appendix C: Jump distribution for the coordination game

We obtain the jump distribution 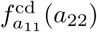 from mutants who playing the coordination game, and compare it to the jump distribution from dominant mutants. The jump distribution for the coordination game can be obtained from the same form of Eq. (14) with a different integrating range

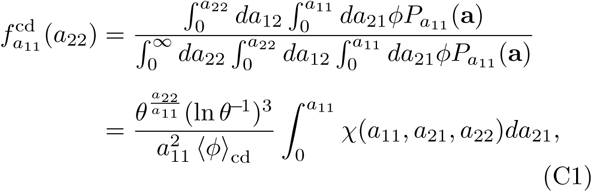

where 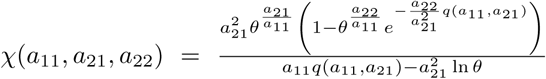 with *q*(*a*_11_*, a*_21_) = *M* (*a*_11_ − *a*_21_)(*a*_11_λ + *a*_21_λ_*d*_). Due to the form of *ϕ* for the coordination game, the integration is more complex than dominance game case. We numerically integrate Eq. (C1) and draw the distributions for various parameters together with the jump distribution for the dominance game, see Fig C1. As we can see in the figure, the distribution 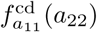 becomes more closer to 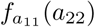 as *a*_11_ becomes smaller. As both distributions become with decreasing *a*_11_, we use the same jump distribution written in Eq. (15) for both games in the main text.

**FIG. C1.**
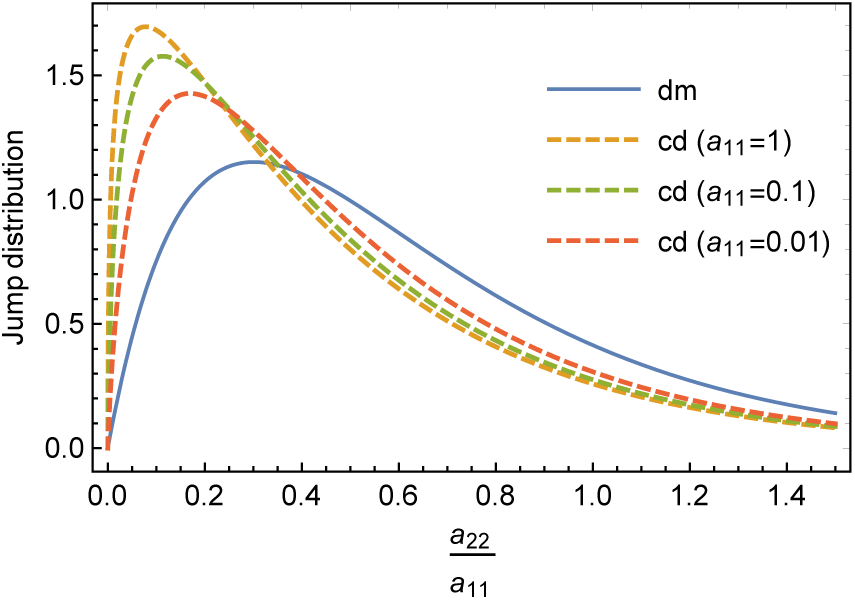
Jump distributions for mutants who playing dominance and coordination games, Eq. (14) and Eq. (C1), respectively. We used *θ* = 0.1 for different *a*_11_ values: *a*_11_ = 1, *a*_11_ = 0.1, and *a*_11_ = 0.01. For the rescalied variable 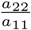, the jump distributions for the dominance game are collapsed in one curve. For the coordination game, the distribution becomes more closer to the jump distribution of the dominance game as *a*_11_ decreases. We used *M* = 100, λ_*d*_ = 0.4, and λ_*b*_ = 0.9.

### Appendix D: Threshold payoff *ã*

Single-type populations with a small payoff *a*_11_ are prone to go extinct due to stochastic fluctuations of the population size. We define the threshold payoff *ã* as a payoff at which the system typically goes extinct before a change of the carrying capacity. Hence, at *ã*, the mean time to extinction of single-type populations is equal to the mean time to the next successful fixation of mutants. Note that we deal with real time *T* (continuous) instead of mutant event time *t* (discontinuous). By simulating an ensemble of populations, we numerically compute the characteristic time to extinction *T*_ext_ of single-type populations without mutation (see Fig. D1). The results of stochastic simulations show a clear exponential pattern. By fitting we obtained *T*_ext_(*a*_11_) ≈ 5 exp(165*a*_11_) at a given parameter set, see Fig. D1. A successful invading mutant appears in *T*_mut_ = *ξ*/*K*λ_*b*_*μ* where *K* = *a*_11_*M*λ. If *T*_ext_ is smaller than *T*_mut_, populations typically go to extinction before the next chance to rescue their population. Hence, the threshold value *ã* can be evaluated by equating both time scales, *T*_ext_ = *T*_mut_. If we assume that only dominant mutants contribute to *ξ*, we can obtain the analytical expression of *ã*,

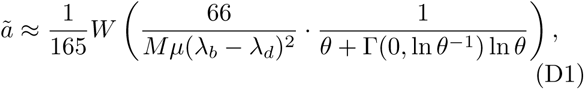

where *W* (*z*) is the Lambert-W function which gives the solution for *w* in *z* = *we*^*w*^.

Figure. D2 shows *ã* in *θ* with and without the coordination game in *ξ*. We numerically solving *T*_ext_ = *T*_mut_ and get the results with coordination game. The order is the same for both cases,𝒪(10^−2^). Despite the discrepancy between two results for *ã*, it does not significantly affect the mean time to extinction (see Fig D2 (b)). Therefore, we use *ã* as expressed in Eq. (D1) for the main text.

### Appendix E: Asymptotic behavior of the average carrying capacity

To calculate the carrying capacity changes in the long run, we need the distribution of payoff *a*_11_ in time. The solution (17) provides us the distribution of *u* = ln(*a*_11_), which leads to

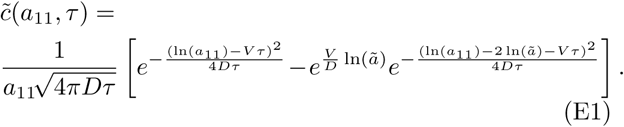

Note that the pre-factor 1/*a*_11_ originates from the variable change. Then, the expectation value of the carrying capacity is

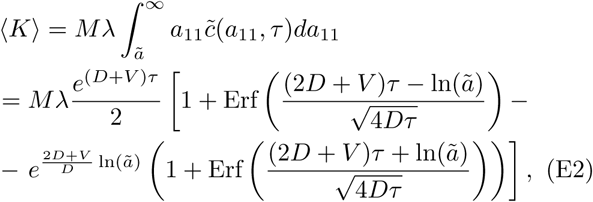

where Erf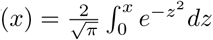.

**FIG. D1.**
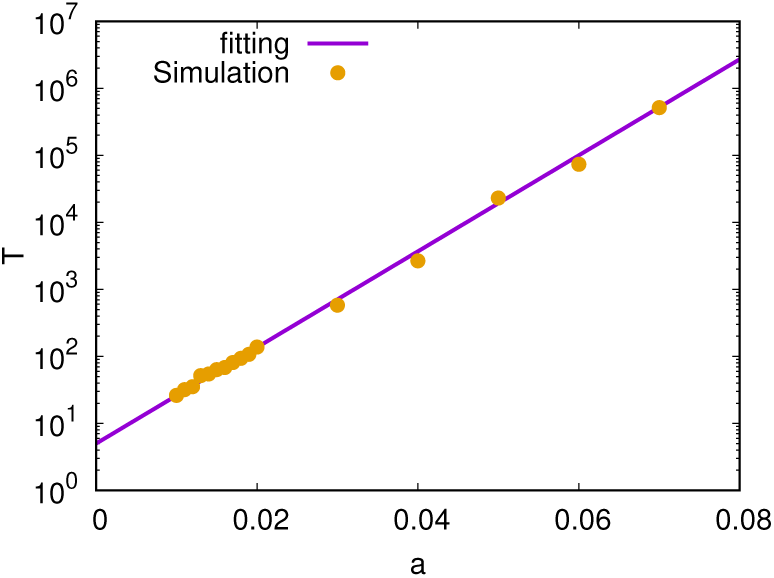
Mean time to extinction *T*_ext_ measured in real time unit for single-type populations without mutation. As we expected, *T*_ext_ increases in *a*_11_ with an exponent 165. The fitting function *f* (*a*_11_) is *f* (*a*_11_) = 5 exp(165*a*_11_) for *M* = 1000. For all simulations, we use λ_*b*_ = 0.9 and λ_*d*_ = 0.4. Since the time scale for a new mutant is *ξ*/*K*λ_*b*_*μ*, most populations go to extinct when *T* < *ξ*/*K*λ_*b*_*μ*. Hence, we define the threshold payoff *ã* by equating two time scales *ξ*/*a*11*Mμ* and 5 exp(165*a*_11_). For the mutation rate, we used *μ* = 10^−5^.

To describe the long time behavior (*τ* → ∞), we use the common approximations

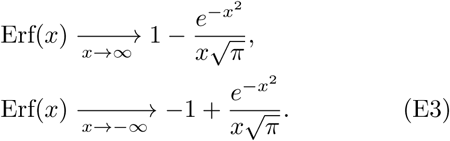

Hence, we get

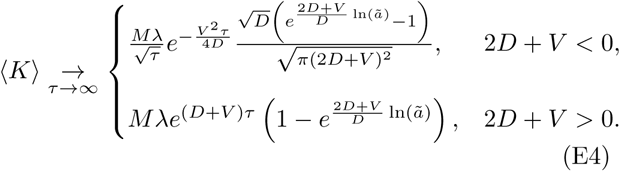

The expected carrying capacity of surviving populations is given by

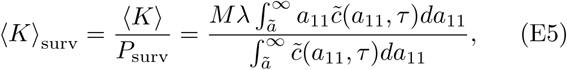

where

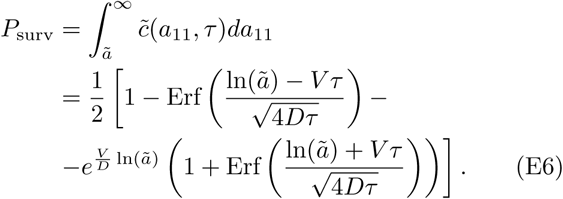

**FIG. D2.**
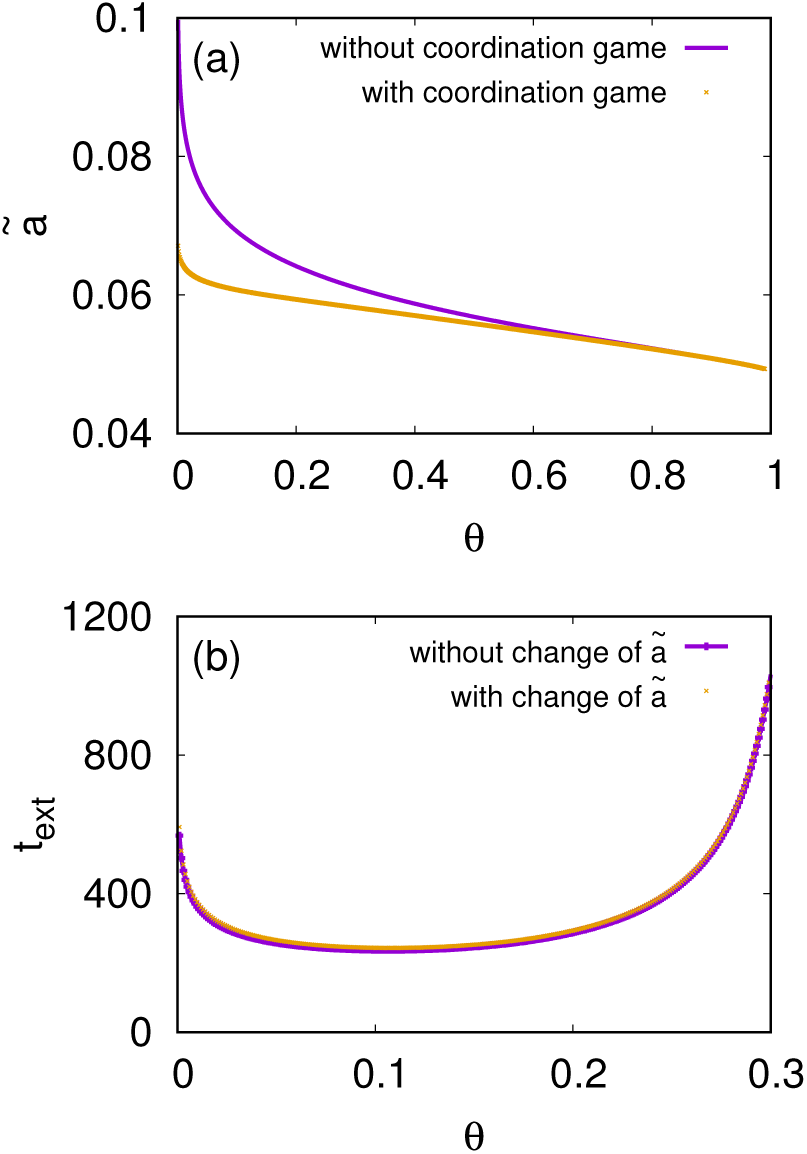
Threshold payoffs decreases in *θ*, and the order of magnitude is 10^−2^. The time scale *ξ* can be calculated from dominance and coordination games. We examine the effect of coordination game on *ã*. As shows in (a), the change of *ξ* from the coordination game does not change order of *ã*. As we can see in (b), the changes of *ã* by the coordination game is not crucial for the results of *t*_ext_. Hence, we keep the expression of *ã* from only taking into account dominance game. We used *M* = 1000, λ_*b*_ = 0.9, λ_*d*_ = 0.4, and *μ* = 10^−5^.

Using Eq. (E3), we get

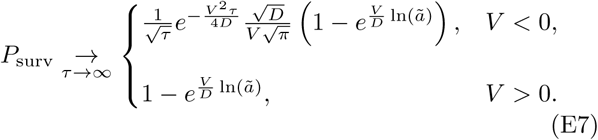

Substituting Eqs. (E4) and (E7) into Eq. (E5), we get 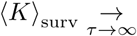

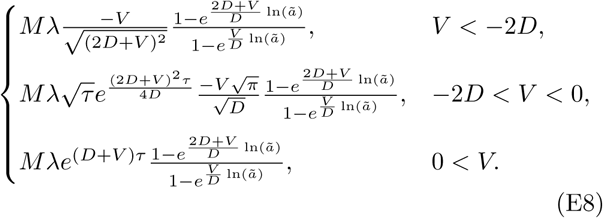

Focussing on the time dependence, we obtain

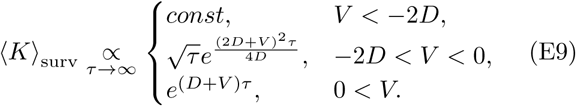

The borders between regimes are at *V* = −2*D* and *V* = 0. These correspond to 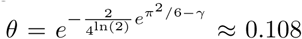 and 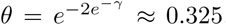, respectively. The border between exponential decline and exponential growth of ⟨*K*⟩ is achieved at *V* = −*D*, equivalent to 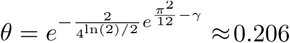.

### Appendix F: Out of weak mutation regime

If the next successful mutant emerges before its maternal population reaches its carrying capacity, the weak mutation assumption is violated. We call this regime the diluted regime, because the population size *N* is much smaller than its carrying capacity *K*. Hence, the competition may not play a major role in this regime. Here, we estimate the population size *N*_*c*_ and the time *τ*_*c*_ to enter the diluted regime. Usually, populations grow to large sizes when *V* > 0, and thus we only focus on *V* > 0. In this parameter range, the coordination game is negligible. Thus we only consider the dominance game for *ξ*.

The average time interval between two successful fixation of mutants is *T*_mut_ in real time unit. On the other hand, the time that mutants reach its carrying capacity can be estimated as *T*_growth_ ≈ ln(*N*)/λ. The population enters the diluted regime when *T*_mut_ ≤ *T*_growth_. This happens at the population size 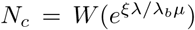. We can obtain the necessary time *τ*_*c*_ to reach the population size *N*_*c*_ from the Eq. (E4),

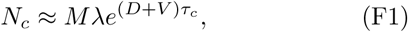

where we neglected the term 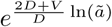. Hence,

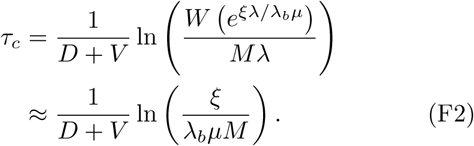

**FIG. F1.**
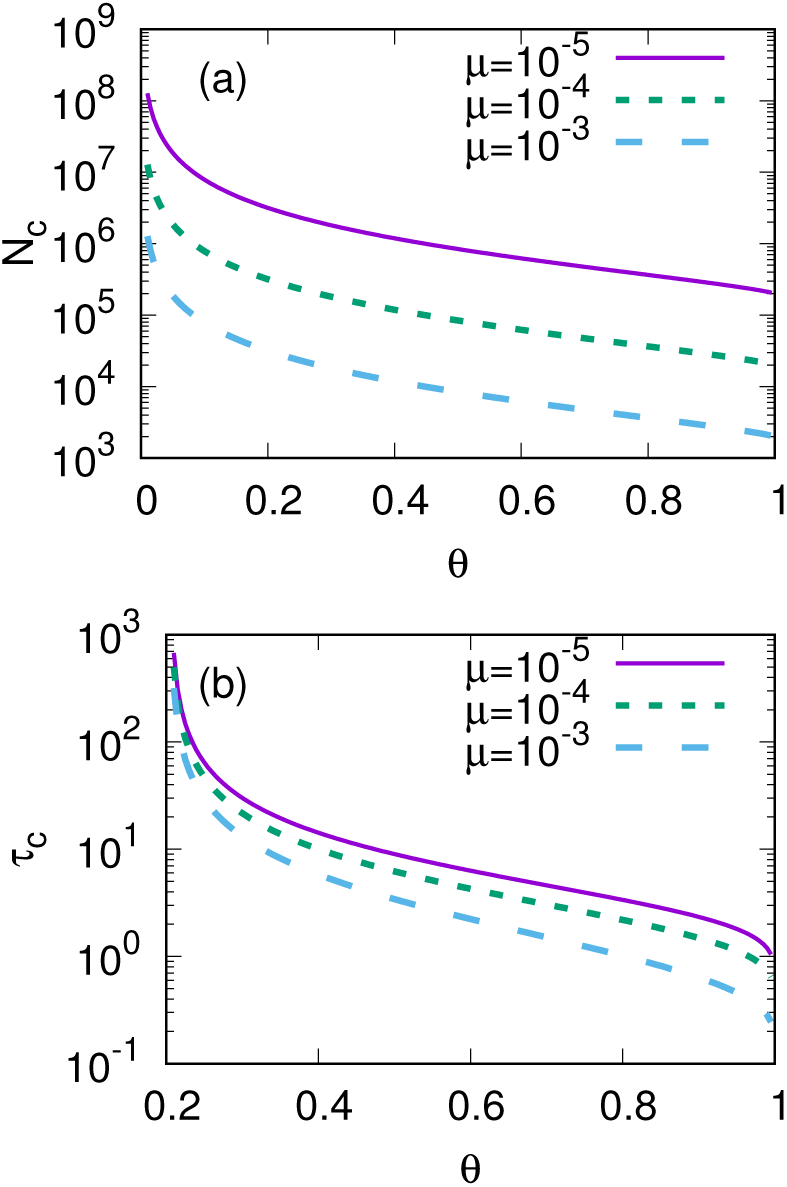
The criteria of population sizes *N*_*c*_ in which the population enter the dilute regime is shown in (a). The time *τ*_*c*_ to reach this population size is drawn in (b). The average population size decreases for *θ* < 0.206, and thus the population does not reach *N*_*c*_ unless otherwise they start from *N* ≥ *N*_*c*_. Hence, we only show *τ*_*c*_ for *θ* > 0.206. Both *N*_*c*_ and *τ*_*c*_ are decreasing in *θ*. We used *M* = 1000, λ_*b*_ = 0.9, λ_*d*_ = 0.4, and *μ* = 10^−5^ for calculations.

In the above approximation, we used *W* (*x*) | _*x* →∞_ = ln(*x*) − ln(ln(*x*)).

In this regime, the population size is far from the carrying capacity, and death from competition does not play an important role. Hence, the constant death and birth rates are determining the population size *N*,

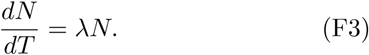

The mutation event time per real time is

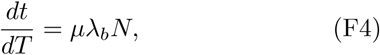

where λ_*b*_*N* gives the number of divisions per real time unit. Combining above two equations, we can obtain

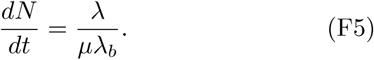

Thus, the population size *N* is linearly increasing in *t*

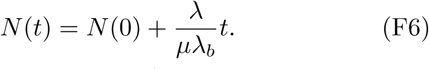

A new mutant emerges after 1/*μ* births, and during one birth, a death happens with probability λ_*d*_*/*λ_*b*_. In a unit time of one birth, the population size changes 1 − λ_*d*_*/*λ_*b*_. Hence, following every mutation event, the population size linearly increases by 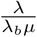. The results also agree well with our prediction, as shown in Fig. 4. The results show a linear growth of population size even when the population size is smaller than *N*_*c*_. This implies that the growth of populations slows down in the long time regime for large *θ*.

## ACKNOWLEDGMENTS

We thank Su-Chan Park, Hyeong-Chai Jeong, and Seung Ki Baek for constructive comments and discussions.

